# MTrack: Automated Detection, Tracking, and Analysis of Dynamic Microtubules

**DOI:** 10.1101/368191

**Authors:** Varun Kapoor, William G. Hirst, Christoph Hentschel, Stephan Preibisch, Simone Reber

**Affiliations:** Imaging Lab, Berlin Institute for Medical Systems Biology (BIMSB), Max Delbrück Center for Molecular Medicine in the Helmholtz Association (MDC), 13125 Berlin, Germany; Quantitative Biology Lab, IRI Life Sciences, Humboldt-Universität zu Berlin, 10115 Berlin, Germany

**Author notes:** **For correspondence:** (SP); (SR). **Present address:** Institut Curie, 75248 Paris Cedex 05, France (Varun Kapoor).

## Abstract

Microtubules are polar, dynamic filaments fundamental to many cellular processes. In vitro reconstitution approaches with purified tubulin are essential to elucidate different aspects of microtubule behavior. To date, deriving data from fluorescence microscopy images by manually creating and analyzing kymographs is still commonplace. Here, we present MTrack, implemented as a plug-in for the open-source platform Fiji, which automatically identifies and tracks dynamic microtubules with sub-pixel resolution using advanced objection recognition. MTrack provides automatic data interpretation yielding relevant parameters of microtubule dynamic instability together with population statistics. The application of our software produces unbiased and comparable quantitative datasets in a fully automated fashion. This helps the experimentalist to achieve higher reproducibility at higher throughput on a user-friendly platform. We use simulated data and real data to benchmark our algorithm and show that it reliably detects, tracks, and analyzes dynamic microtubules and achieves sub-pixel precision even at low signal-to-noise ratios.

## Introduction

Microtubules are dynamic filaments essential for many cellular processes such as intracellular transport, cell motility and chromosome segregation. They assemble from dimeric *αβ*-tubulin subunits that polymerize in a head-to-tail fashion into polar filaments [26] (Figure 1). Microtubules show a behavior termed ‘dynamic instability’, which can be empirically described by four parameters: (1) the polymerization velocity at which microtubules grow (*vg*), (2) the depolymerization velocity at which microtubules shrink (*vs*), (3) the catastrophe frequency at which microtubules switch from growth to shrinkage (*fc*), and (4) the rescue frequency at which microtubules switch from shrinkage to growth (*fs*) [25]. This dynamic behavior is intrinsic to microtubules. In a cellular context, however, the dynamic properties of microtubules are modulated by motors and accessory proteins known as microtubule associated proteins (MAPs) [6, 44, 5, 31, 10]. In most cases, the cellular context is too complex to study a single protein’s contribution to microtubule dynamics. Therefore, biochemical activities of individual proteins have primarily been characterized in vitro using purified components and total-internal reflection fluorescence (TIRF) microscopy [32, 37, 15, 24, 7, 12, 39, 3, 13, 46]. Furthermore, microtubule dynamics are strongly affected by a set of drugs routinely used to treat diseases such as cancer [17] and malaria [18]. Owing to their clinical relevance, it is a viable need to understand the exact regulation of microtubule dynamics by a given drug and thereby elucidate the underlying molecular mechanisms. Given the growing interest in biochemical reconstitution systems [6, 44, 9], automation of data analysis will unveil the full potential of the experimental approaches as described above.

**Figure 1.**
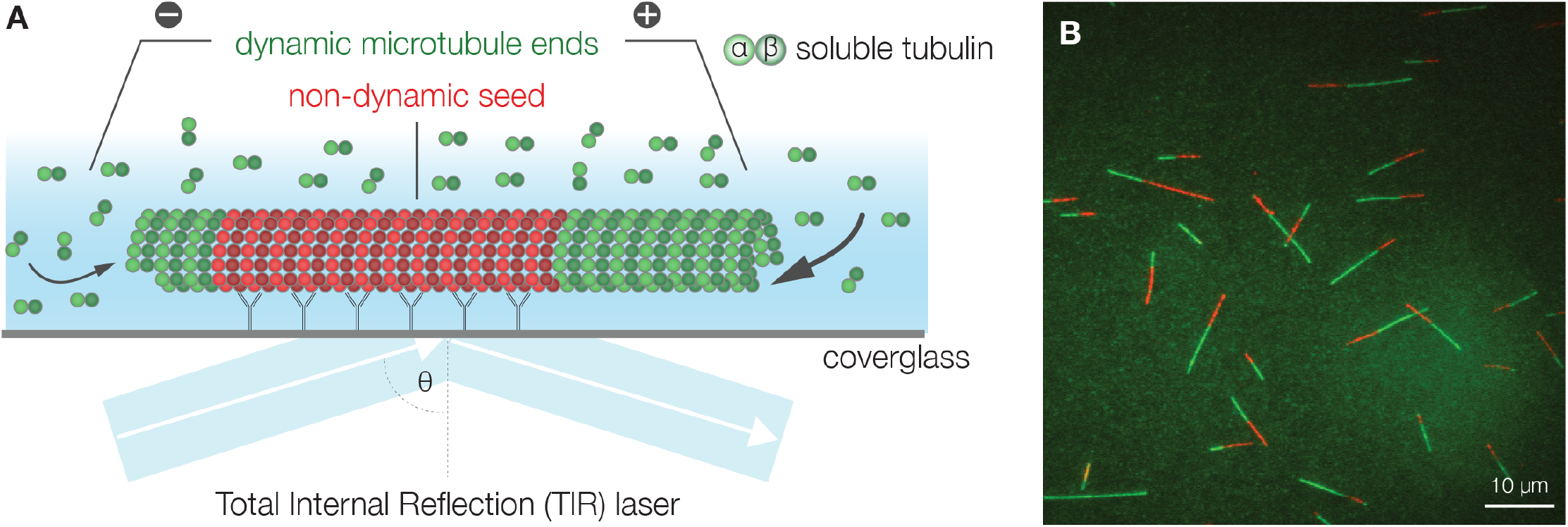
Microtubule Dynamics by TIRF Microscopy. **A** Schematic experimental design: Stabilized microtubule seeds (red) are bound to the coverglass by antibodies and serve as nucleation points for dynamic microtubules (green). One microtubule end usually shows higher growth rates (+) than the other end (−). Total internal reflection fluorescence (TIRF) microscopy selectively excites fluorophores in a restricted volume adjacent to the glass-water interface allowing the visualization of individual microtubules. **B** TIRF microscopy image of dynamic microtubules (green) grown from stabilized seeds (red).

Quantitatively deriving dynamic microtubule parameters from fluorescence microscopy images by manually creating and analyzing kymographs (spatial position over time) is still common practice [49]. This limits the collection of statistically significant amounts of data. Moreover, manual analysis can bias data collection and introduce variability. Thus, methods have been developed that allow microtubule detection and/or tracking [48, 4, 23, 30, 8, 34]. However, to date, there is no fully automated workflow that provides detection and tracking of microtubules followed by automated data analysis and statistics collection. Here, we present the software MTrack, which detects, tracks, measures, and analyses the behavior of fluorescently labeled microtubules imaged by TIRF microscopy (Supplementary Figure 1). MTrack is capable of automatically identifying and tracking dynamically growing microtubules that potentially bend and cross with subpixel resolution, even at high growth rates and low signal-to-noise ratios (SNR) using advanced objection recognition and robust outlier removal. The software is easily accessible for users and developers since it can be automated and is provided as an open-source Fiji [36] plug-in.

## Results

The MTrack software is organized in two consecutive modules that can be run independently. The first module robustly detects microtubule seeds and tracks dynamic microtubules over time. The second module interprets the length over time plots to extract relevant parameters of dynamic instability and population statistics.

### Robust Detection of Microtubule Seeds using MSER and Sum of 2D Gaussians

A common way to reconstitute and analyze microtubule dynamics is by the use of TIRF microscopy and fluorescently labeled tubulin [39, 13]. Stabilized (non-dynamic) fluorescent microtubule seeds are immobilized onto glass surfaces. These microtubule seeds serve as nucleation points from which dynamic fluorescent microtubules will grow (Figure 1). Previous microtubule tracking software relies on manually clicking each individual microtubule to be analyzed [4, 34]. Therefore, our aim was to develop an approach that robustly detects microtubule seeds in the image in a fully automated fashion. It is essential to precisely determine the exact end point of each seed, as these are the sites from which microtubules will subsequently grow and shrink. MTrack does so by using the Maximally Stable Extremal Regions (MSER) algorithm [21, 28] to identify image areas belonging to each seed, a sum of 2D Gaussians (SoG) model to accurately localize individual seeds, and finally a Gaussian Mask fit [40] to determine the precise end point of each seed with subpixel resolution (Figure 2A).

**Figure 2.**
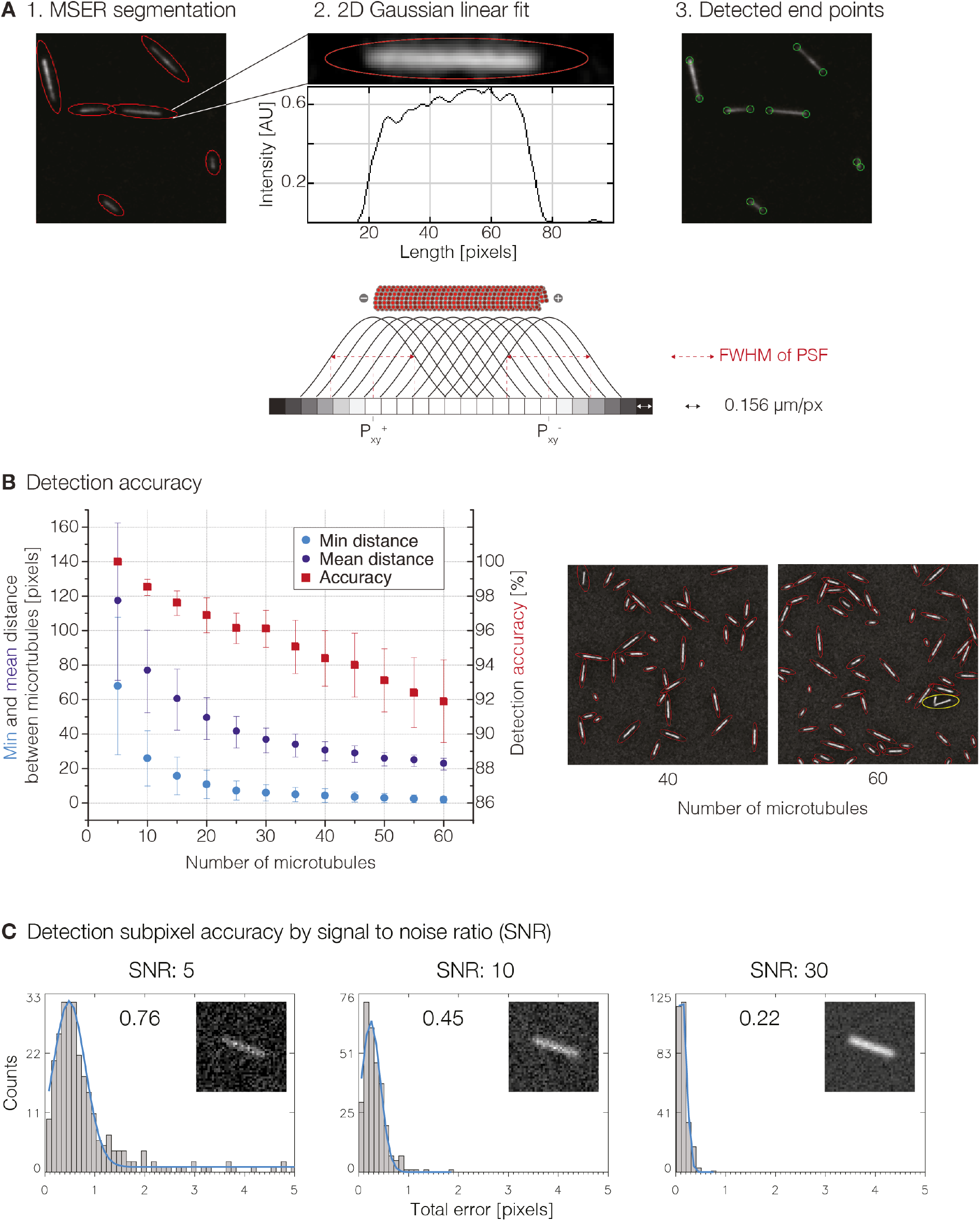
Microtubule Seed Detection. **A** The default algorithm to identify microtubule seeds as objects is Maximally Stable Extremal Regions (MSER). 1. Microtubule seeds detected by MSER are marked by fitted red ellipses. 2. To determine the extact end point of each microtubule seed, a sum of 2D Gaussian model is fit to the major axis of the ellipsoid. FWHM (full width at half maximum) of the PSF (point spread function) is determined experimentally. 3. Detected end points are marked by green circles. **B** Detection accuracy of microtubule seeds with a close to 100% accuracy when distance between seeds is larger than 5 pixels. Detection error is marked in yellow. **C** Subpixel accuracy of microtubule seed end point detection depends on SNR. Values correspond to detection error in pixel.

The principle underlying MSER is a component tree, which computes every possible threshold of the image thereby increasing the dimensionality of the input image by one (e.g. 2d > 3d, Supplementary Movie 1). Stable regions within the component tree are those that do not significantly change over multiple thresholds. Since microtubules can vary in size, are randomly oriented, potentially bent, and are the main bright objects in the fluorescent image, the MSER detector performs accurately without the need to make assumptions about shape, orientation, and size of regions. Successfully detected microtubule seeds show a one-to-one assignment to ellipsoidal regions (Figure 2A). Even for low SNRs, the overall detection accuracy mostly depends on the density of microtubule seeds (Figure 2B). MTrack detects seeds with a close to 100% accuracy when the distance between seeds is larger than 5 pixels, which is experimentally feasible. Using the region identified by MSER, we fit a SoG model (see Supplementary Material) using the major axis of the ellipsoid as a starting point to detect the accurate end position of the seeds (Figure 2A). Final end points are computed using a modified Gaussian Mask fit [40] to maximize location precision (see Supplementary Material). It uses combined Gaussian distributions to model the appearance of the end points in the image as defined by their location and the point-spread function of the system (Figure 2A).

The signal-to-noise ratio (SNR) is directly related to the maximally achievable localization precision [40, 4]. Therefore, we simulated microtubule seeds with different SNRs and show that at reasonable experimental SNRs (Material and Methods, Supplementary Figure 2) endpoints can be accurately localized with subpixel resolution (Figure 2C). Despite the fact that microtubule ends show no symmetric intensity distribution, the detection error is normally distributed showing no bias towards microtubules pointing in either direction (Supplementary Figure 2). Therefore, detection precision is robust and does not depend on filament orientation.

### Tracking Microtubules by 2D Gaussian Polynomial Models

The goal of microtubule tracking is to identify the end points of growing and shrinking microtubules within each frame of a fluorescent time-lapse movie with the highest possible accuracy and reliability. Typically such data is displayed as a length versus time graph (kymograph). Microtubules stochastically switch between growth and shrinkage. Growth velocities observed in vitro range from 0.6 μm/min [45] to 40 μm/min [47]. Moreover, depolymerization velocity is typically an order of magnitude greater than the corresponding growth velocity [49]. Thus, the frame-to-frame difference in microtubule length can be significant. In addition, brightness changes, growth heterogeneity, microtubules bending, crossing, and moving out of focus present further challenges to accurate tracking. We will show that fitting polynomial functions enables us to robustly track microtubules, even when bending or crossing.

In more detail, MSER first detects an image region for each dynamic microtubule within each frame of the fluorescent time-lapse movie (as described for seed detection). To initialize the iterative microtubule detection within each MSER region, we need a start point and the guess of an end point. The start point is fixed and defined by the detected seed end, while the end point is estimated by the intersection of the current MSER region boundary with the projected growth direction from the last successfully segmented frame, which can potentially be many frames away (Figure 3A, see also discussion). These two points define a line used to initialize the fitting of a 2D SoG model to the image region that identifies the microtubule. However, the 2D SoG path is represented by a 3^*rd*^ order polynomial function, which enables tracking of bending (Figure 3B) and crossing microtubules (Figure 3C, Supplementary Material). The iterative fitting of the 2D SoG model is robust and will recover even after many frames have been missed since it is able to identify the correct microtubule path as long as the initialization line intersects with the microtubule at any point. Additionally, we reject estimates of microtubule paths in the current frame that differ significantly from the previous time-point (see also discussion, Supplementary Material). The precise endpoint of the growing or shrinking microtubule is finally computed using our modified Gaussian Mask fit as described for seed detection.

**Figure 3.**
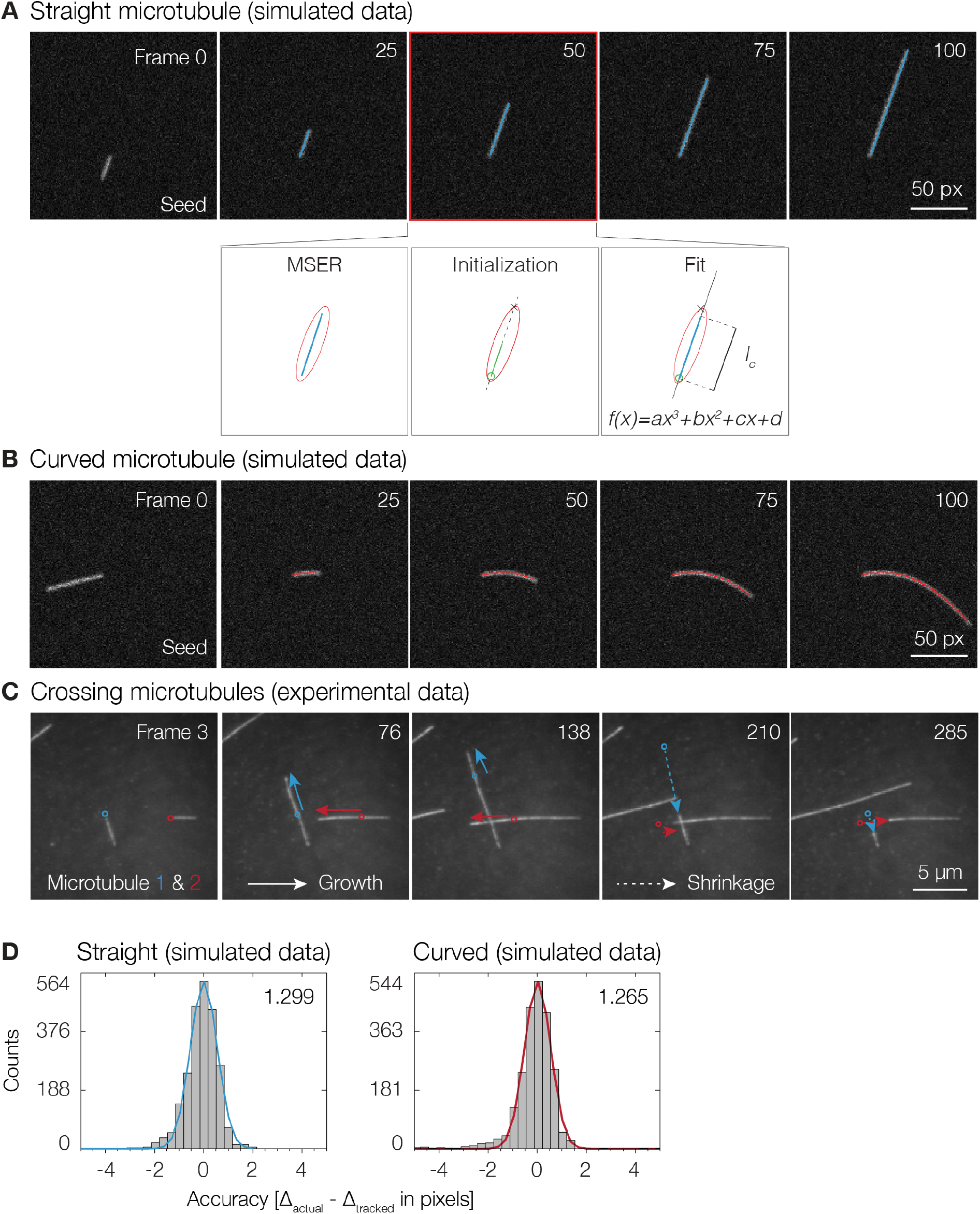
Microtubule Tracking. The MTrack algorithm successfully tracks straight, bending and crossing microtubules over time. **A** First, MSER detects an image region (red ellipse) for each dynamic microtubule (blue). The seed end is then identified as a start point (green circle) and the end point (x) is estimated by the intersection of the current MSER region boundary with the projected growth direction (dashed line) from the last successfully segmented microtubule. These two points initialize a 2D SoG fit represented by a 3^*rd*^ order polynomial function. The actual length of the dynamic microtubule is calculated as the contour length (*lc*) of the final fit. This approach allows tracking of bending **B** and crossing **C** microtubules. **D** Tracking accuracy for straight and bending microtubules (SNR 10) was determined as the distance between the actual simulated position of the microtubule end (Δ_actual_) and the position given by the tracking algorithm (Δ_tracked_).

When imaging dynamic microtubules, background fluorescence is inevitable as fluorescent tubulin in solution is required to allow microtubules to grow. With high tubulin concentrations, background fluorescence increases and therefore the “effective SNR” of the microtubule signal considering the fluorescent background can be considerably low (Supplementary Figure 2). To characterize tracking performance, we simulated sequences of dynamic microtubules assuming different SNRs. Such an approach was previously demonstrated to be valuable for the study of the localization precision in single molecule localization microscopy [38] and found to be a good single predictor of tracking precision [4]. Our analysis shows that irrespective of growth along a line or a 3^*rd*^ order polynomial, the error distribution remains similar and achieves pixel resolution (Figure 3D).

### Automatic Derivation of Microtubule Dynamic Parameters using Iterative Robust Outlier Removals

The aim of this module is to extract relevant microtubule dynamic parameters from length versus time graphs (Figure 4A). Each microtubule track has an unknown number of growth and shrinkage events. To automatically derive these dynamic parameters, we developed an iterative, model-based robust outlier removal algorithm based on RANSAC (Random Sample Consensus) [11]. RANSAC is a non-deterministic algorithm that fits a function to data points by maximizing the number of data points that support the function fit (inliers), given an error *є* (Figure 4B).

**Figure 4.**
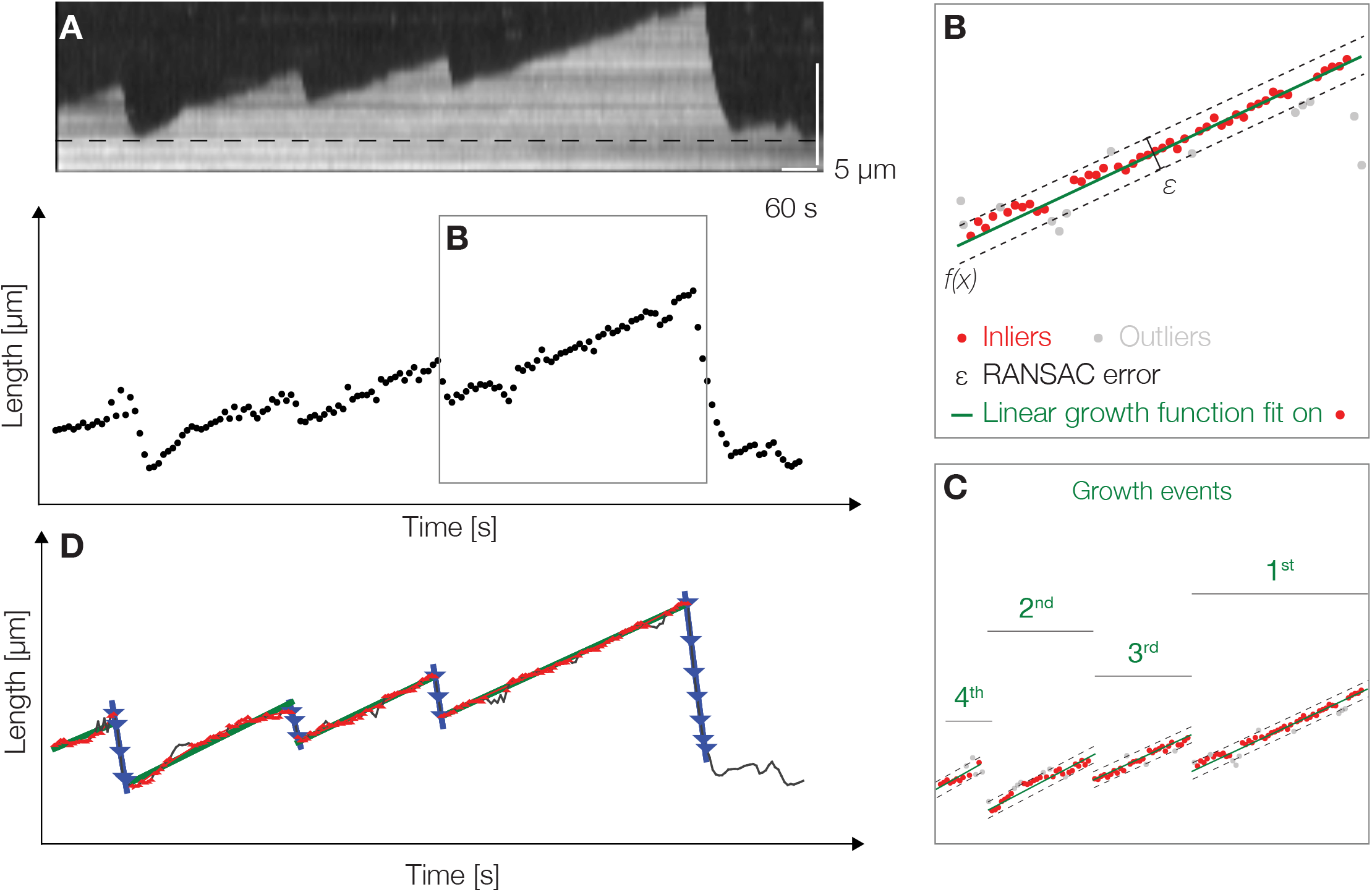
Automated Derivation of Parameters of Microtubule Dynamic Instability using Iterative Robust Outlier Removal. **A** Kymograph of a dynamic microtubule. **B** In the length versus time plot obtained from tracking, RANSAC first identifies the largest subset of consecutive time points that follow near-linear growth as a growth event. **C** To identify all growth events, RANSAC iteratively removes time points belonging to an identified growth event from the sampling set and repeats the RANSAC sampling until no further growth events can be found. **D** The final graph containing detected growth and shrinkage events, as well as catastrophes and rescues.

**Figure 5.**
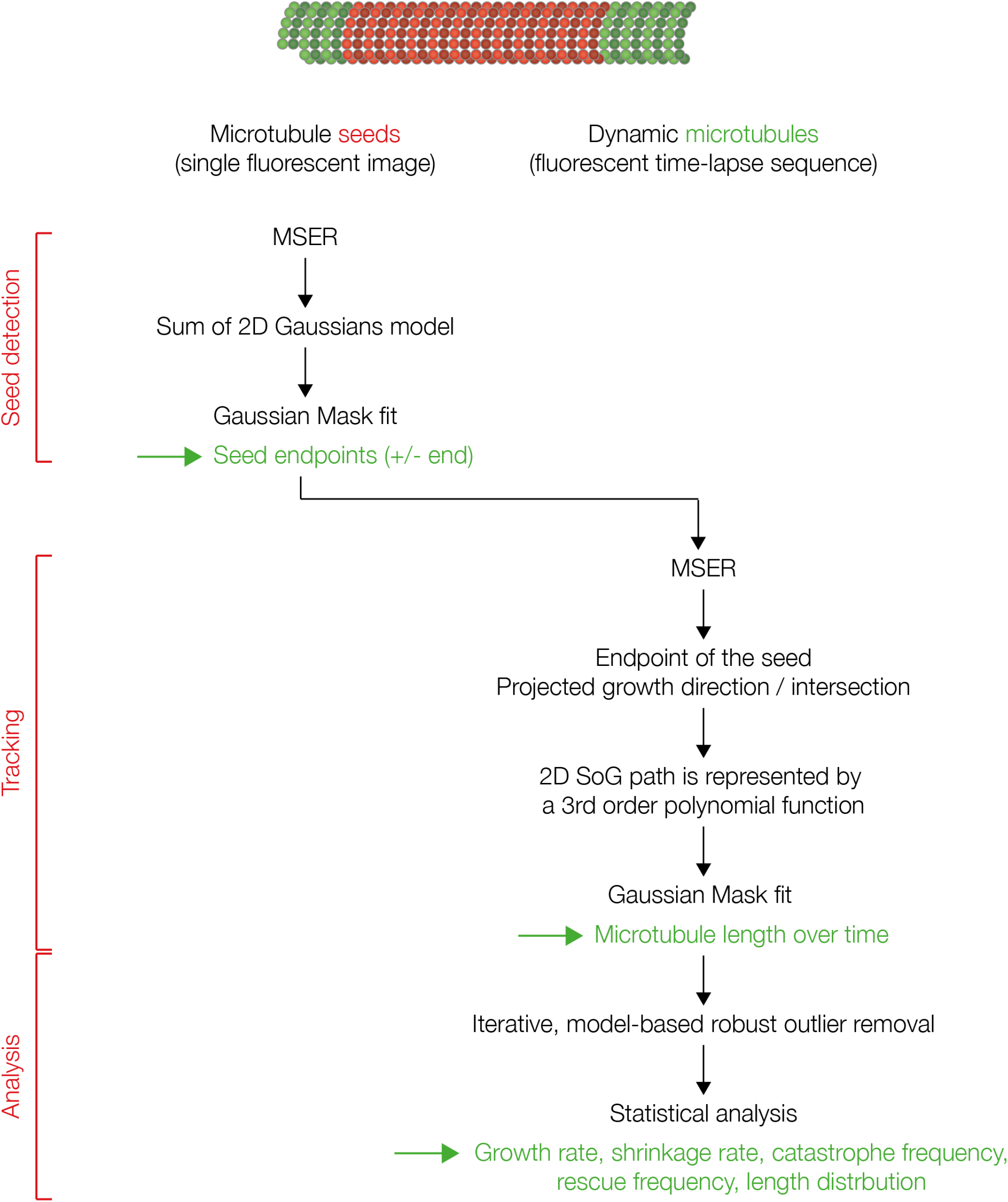
Flow Diagram of MTrack Detecting, Tracking, and Analyzing Dynamic Microtubules

**Figure 6.**
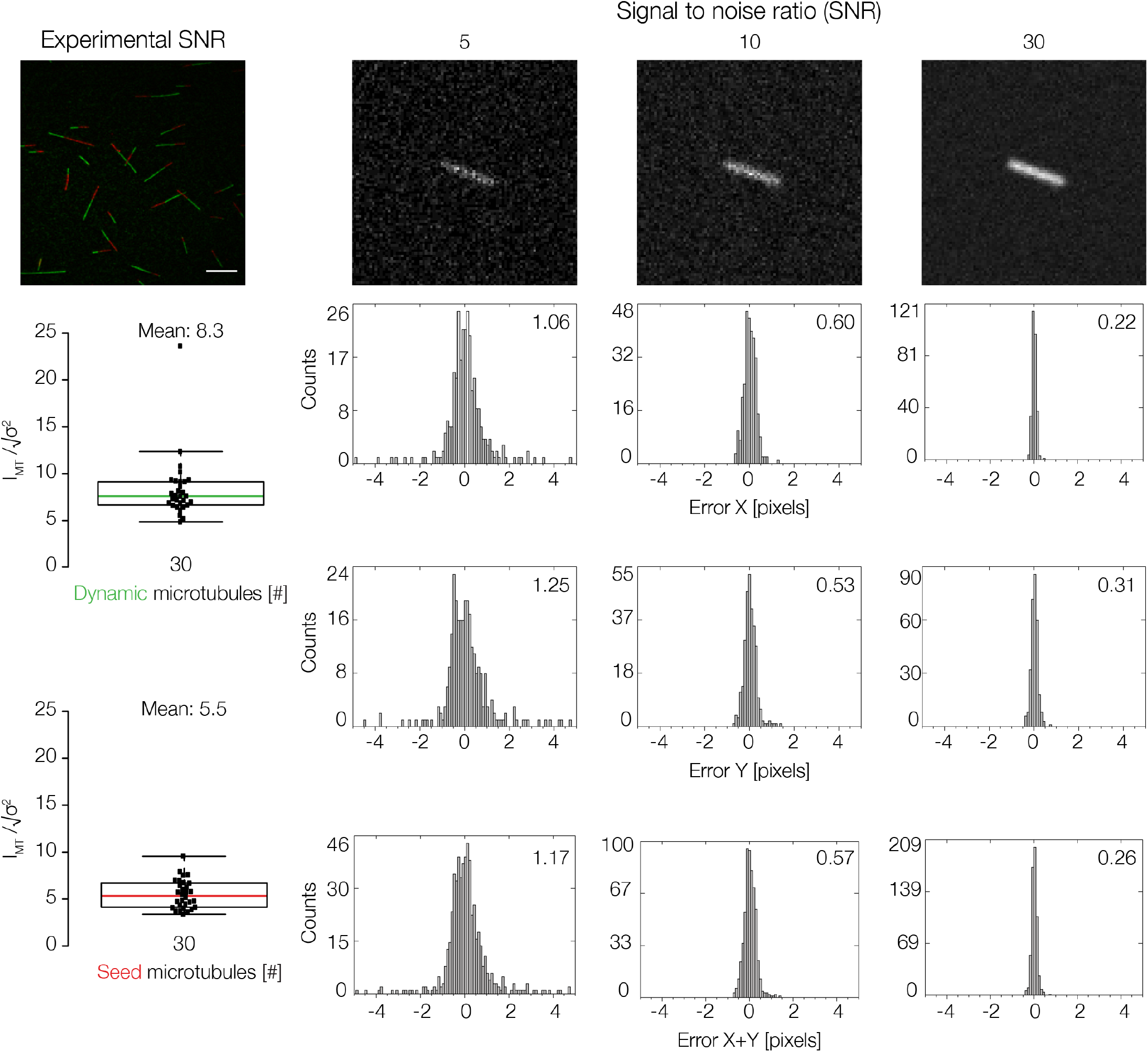
Microtubule Seed Detection. The experimental SNR of dynamic growing microtubules and seeds have been measured and calculated as described in Material and Methods. Scale bar: 10 μm. Subpixel accuracy of simulated microtubule seed end point detection dependens on SNR. Detection error is normally distributed showing no bias towards microtubules growing in either direction.

**Figure 7.**
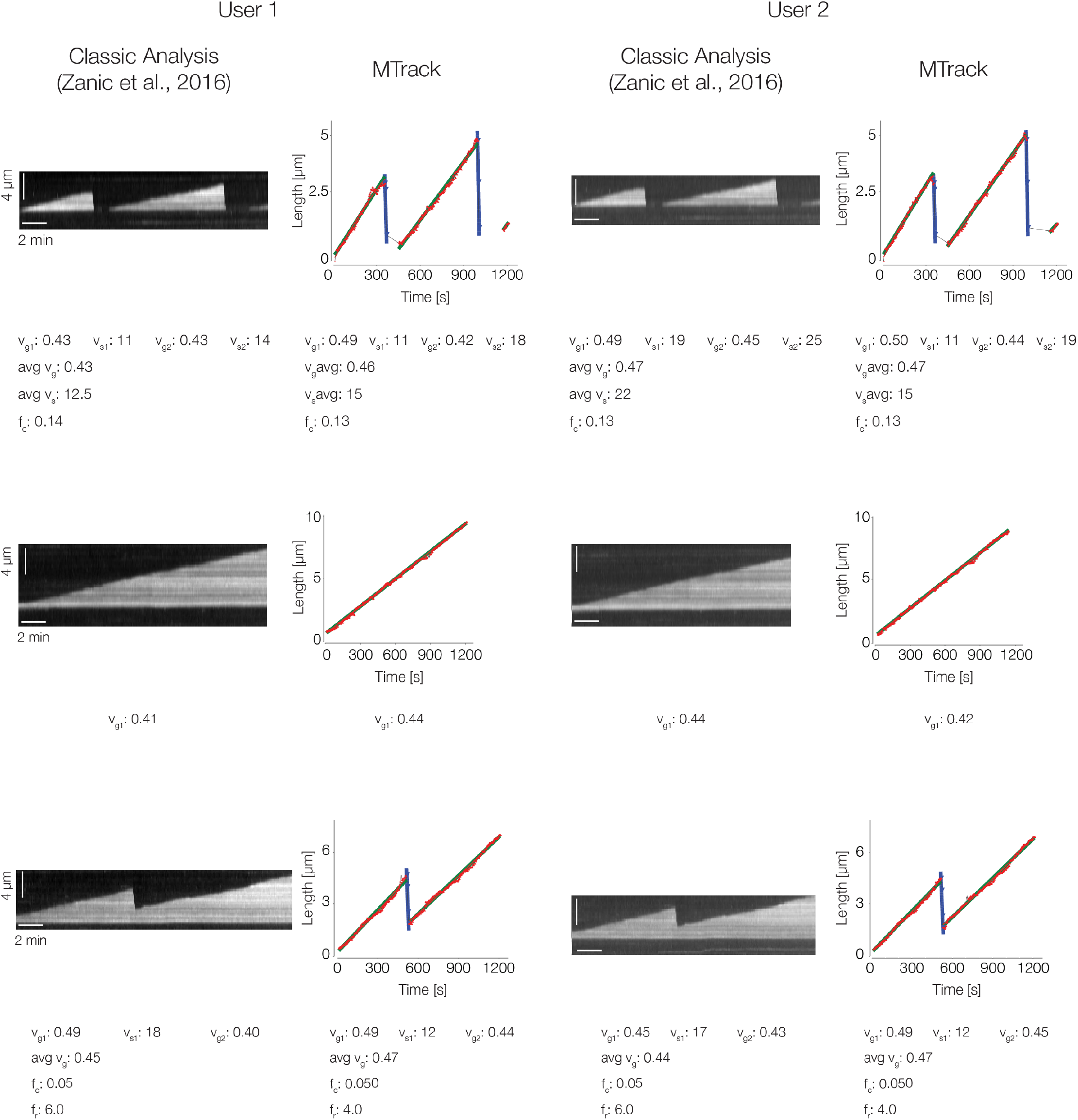
Comparison between Manually Analyzed and Computer-inferred Microtubule Dynamics. Polymerization velocity (*vg*) and depolymerization velocity (*vs*) are given in μm/min, catastrophe frequency (*fc*) and rescue frequency (*fs*) in s^-1^.

First, all growth events are detected. Microtubule polymerization velocity (*vg* [nm/s]) was shown to be a linear function of tubulin concentration and is usually approximated by a single constant growth rate [12]. We therefore assume that periods of microtubule growth follow an increasing, near-linear polynomial function, which is implemented as a 2^*n*^ *d* or 3*^r^d* order polynomial regularized with linear function. To do so, RANSAC first identifies the largest subset of consecutive time points that follow near-linear growth (Figure 4B). To identify all growth events in one track, the software iteratively removes time points belonging to an identified growth event from the sampling set. It then repeats the RANSAC sampling until no further growth events can be found. In the example shown in Figure 4, RANSAC identifies four growth events in order of their length. To achieve compatibility with the general accepted assumption of linear microtubule growth, the software finally fits a linear function to all inlier points of each growth event (Figure 4C). Next, RANSAC identifies events of microtubule shrinkage. Microtubule depolymerization velocity (*vs* [nm/s]) can be an order of magnitude higher than polymerization velocity and therefore, depolymerization events tend to be short. We therefore use a linear model limited to fast decline for our iterative RANSAC algorithm (Figure 4D, blue lines). Subsequently, catastrophe frequency and rescue frequency are calculated. Catastrophe frequency (*fc* [s^-1^]) is determined by dividing the total number of identified shrinkage events by the total time the microtubules were growing (events/time), taking only full growth events into account. Analogously, rescue frequency (*fr* [s^-1^]) is determined by dividing the total number of identified growth events by the total time the microtubule spent shrinking (events/time). We distinguish between ‘total’ catastrophes, when the microtubule shrinks all the way back to the seed, and rescues by comparing the start of the new growth event to the baseline, which is given by the end point of the seed (Material and Methods). As mentioned in the introduction, microtubules have an intrinsic polarity. The microtubule plus end typically grows significantly faster than the microtubule minus end [26]. When both microtubule ends are selected and tracked, MTrack automatically assigns the faster growing end as the plus end. Finally, given the four parameters of dynamic instability, the microtubule length distribution can be computed (based on [43]) at a user chosen time-point or averaged over all the time points to obtain a time-averaged length distribution.

## Discussion

In the last decade, advances in high-resolution techniques have shed new light on how the dynamic behavior of microtubules is modulated (as reviewed in [14]). Here, we present new software, MTrack, which automatically identifies and tracks dynamic microtubules with subpixel resolution and provides automatic data interpretation and statistics. MTrack will be valuable to any experimentalist (1) who studies different microtubule populations [42, 41], (2) who aims to characterize the effect of a MAP or motor protein on microtubule dynamics [32, 37, 15, 24, 7, 12, 46, 3] or (3) who is interested in characterizing the molecular mechanism of microtubule binding drugs [41]. We believe that MTrack is a powerful tool, which together with high-resolution imaging will provide unbiased, high-accuracy data with sufficient statistics to help elucidate new mechanisms of microtubule dynamics control.

The demand for software, which automatically detects and tracks microtubules is mirrored by the recent development of several software packages that allow microtubule detection and/or tracking [48, 4, 23, 30, 8, 34]. The most recent approach by Bohner and colleagues is a further development of the previously published tracking software FIESTA [34] by optimizing it for low SNRs. In other words, the former software [34] is ideal to track non-dynamic microtubule seeds, for example in single-motor stepping assays [33], while the latter [4] is well-suited for tracking microtubules in dynamic growth assays with free fluorescent tubulin. In addition, in order to validate the tracking performance across a broad range of conditions, Bohner and colleagues tracked simulated dynamic microtubules with varying experimental conditions. While they found the fluorophore labeling density, the pixel size of the image as well as the exposure time to be important parameters, they found the SNR a good single predictor of tracking precision [4]. Motivated by this observation, we systematically varied the SNR ratio and determined the detection and tracking accuracy of our software. In Module 1, endpoints of non-dynamic microtubule seeds can be accurately localized with subpixel resolution even at low SNRs. Moreover, the detection error is normally distributed showing no bias towards microtubule direction. From this we concluded that the detection precision was robust even at experimentally relevant SNRs and did not depend on the filament orientation. We then showed that allowing the final fit to follow a 3^*rd*^ order polynomial function enabled us to track straight, bending, and crossing microtubules. The tracking accuracy achieved pixel resolution, providing a close to molecular precision.

While the above mentioned programs reliably track microtubules with nanoscale precision, both approaches require the manual selection of each individual microtubule to be tracked, neither of the program offers automated data analysis nor do the authors comment on how either software performs on bending or crossing microtubules. MTrack is a fully automated workflow that provides detection and tracking of microtubules and is – to our knowledge – the first software that offers automated data interpretation using iterative robust outlier removal. Directly tracking microtubules and analyzing their dynamics is different to tracking and analyzing end-binding proteins (EBs) [1, 22]. Microtubule end-binding proteins such as EB1 accumulate exclusively at growing microtubule ends [2, 24] but get lost once microtubules shrink. In such datasets, microtubule behavior has to be interpolated during phases of microtubule shortening and pausing. In contrast, MTrack can make use of the full length over time information giving by the tracking module to extract information about MT dynamics.

In sum, MTrack is the first software that reliably detects, tracks, and analyzes the behavior of dynamic microtubules. Each module is automated yet highly adaptable and can be used (1) to robustly detect the end points of any linear, fluorescent, filamentous structure, (2) to reliably track fluorescent structures or (3) to analyze length over time plots in an automated fashion.

## Materials and Methods

### Point Spread Function (PSF)

The Point Spread Function (PSF) of a microscope is well approximated by the images of single subdiffraction-sized fluorescent beads. We quantified the PSF and the resolution by fitting 2D Gaussian functions to individual beads for each wavelength. The estimated resolution of the microscope based on the mean value of the full width half maximum (FWHM) is 199 nm and 205 nm for the 647 nm and the 561 nm laser, respectively.

### Signal-to-noise Ratio (SNR)

After background subtraction, we measure and calculate the SNR of microtubules by *I_MT_* – *σ_MT_* where *I_MT_* is the mean deviation of the pixel intensities along a line scan of a microtubule and *σ_MT_* the standard deviation.

### Experimental Data, TIRF microscopy and Imaging

The in vitro reconstitution of microtubule dynamics was performed using tubulin purified from Xenopus egg extracts as described in [46] as well as bovine brain tubulin purchased from Pur-Solutions. Polymerization reactions were carried out at 37°C in BRB80 buffer containing 80 mM PIPES, 1 mM MgCl_2_ and 1 mM EGTA at pH 6.8 supplemented with 1% *β*-mercaptoethanol, 1 mg/mL casein, and 1 mM GTP. Dynamic microtubules were nucleated on GMPCPP-stabilized bovine tubulin microtubules containing 10% Cy5-labeled and 20% biotin-labeled tubulin assembled according to [13, 39]. Reaction chambers were constructed from glass coverslips and slides passivated with dichlorodimethylsilane [27].

Images were taken on the motorized inverted Nikon Eclipse Ti-E microscope with a motorized TIRF angle controlled with ND acquisition software, equipped with a Nikon Plan Apochromat 100x/1.5NA oil immersion objective lens and an EMCCD, Andor iXon Ultra X3 987 Camera. Cy5-labeled seeds were imaged with a 647 nm laser at 1.5 mW (0.2%) and 500 ms exposure time. Dynamic micro-tubules (by mixing Cy3-labeled tubulin with unlabeled tubulin at a ratio of 1:10) were imaged with a 561 nm laser at 0.54 mW (1.2%) and 500 ms exposure time. Image size is 512×512 with a pixel size of 156 nm.

## Mathematics

### 1. Fitting Microtubule Intensity Models

MTrack fits a microtubule’s pixel intensities by using a model, which is a sum of Gaussians along a polynomial. In order to determine the model parameters, we perform minimization of *∼* squared function defined as 
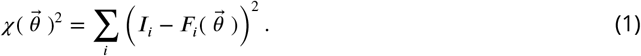

The sum is over the pixel co-ordinates (*i_x_, i_y_*) and *I_i_* is the pixel intensity at that position. *F_i_* represents the value given by the model at the same pixel co-ordinate, and model parameters are represented by the vector 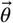. The components of the vector are dependent on the model used (Line model or 3^*rd*^ order polynomial to approximate a beam model [16]). These components will be described in the sections detailing the respective model.

### 2. Line Model

For the seed image, MTrack uses a first order polynomial. A first order polynomial is a line in the image and in this model we place Gaussians along the line. The function *F* introduced before has the following form: 
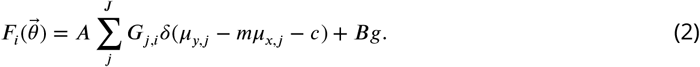

Here, we placed Gaussians enumerated by index *j*. *δ* denotes the dirac-delta function that constrains the location of the Gaussians to be along a function, which is in this case a line. *J* defines the total number of Gaussians in the sum. The centroid of the Gaussian *j* is given by (*μ_x,j_, μ_y,j_*). By using a delta function we ensure that these centroids are always along the line, which is defined as the argument of the delta function. *B_g_* is the background intensity term, *A* is the amplitude of all Gaussians *J*. *G_j,i_* defines a two dimensional Gaussian located at pixel location (*i_x_, i_x_*). In our sum of Gaussian model, the intensity contribution of Gaussian *j* at pixel location *i* is given by 
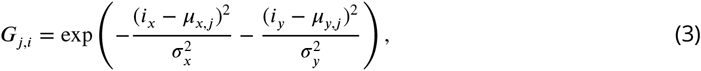

*σ_x_* and *σ_y_* are determined by the point spread function of the microscope. The spacing between the successive Gaussians placed along a curve is assumed to be a constant, defined by the parameter *ds* 
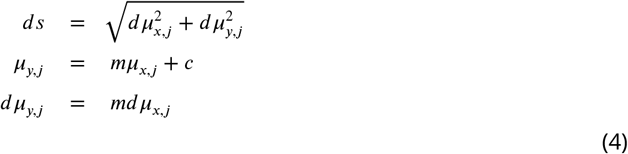

Using these expressions we can write *ds* in the above expression as 
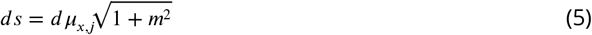

Using the expression above and expanding the expression in 2 we obtain modelled intensities for each pixel *i* by 
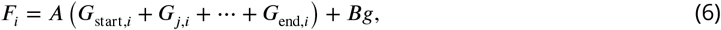

*G*_start,*i*_ and *G*_end,*i*_ define the two dimensional Gaussian at the start and the end location. The modelled intensity of each Gaussian *G_j,i_* along a line is defined as 
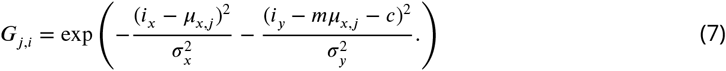

For the seed image the line parameters (*m* and *c*) are determined by using MSER [21]. *σ_x_* and *σ_x_* are user input for the PSF of the microscope and are not fit parameters. The fit parameters to be determined are the start and the end sub-pixel co-ordinate of the model (*μ_x_*,_start_, *μ_y_*,_start_) and (μ*_x_*,_end_, *μ_x_*,_end_) respectively, the spacing between the Gaussians *ds*, the background term *Bg* and the amplitude *A*.

In order to determine these parameters, the function defined in 1 is minimized using the Levenberg-Marquardt solver [19, 20]. Such a minimization requires providing derivatives with respect to the fit parameters. MTrack uses analytical derivatives for that and their form is described in the next section.

### 3. Analytical Derivatives for Line Models

#### 3.1 Derivative with respect to *μ_s_*

Using the expression in 6, we can write the derivative with respect to the first term as 
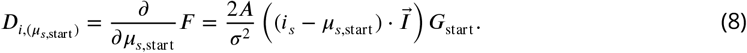

Here, we have used the vector notation to represent the co-ordinates (*s* = 0, 1) for (*x, y*) co-ordinates respectively and 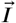 represents the unity vector. The derivative with respect to the end point is also similar to above with *μ_s,start_* being replaced by *μ_s,end_* and *G*_start_ being replaced by *G*_end_.

#### 3.2 Derivative with respect to *ds*

We define 
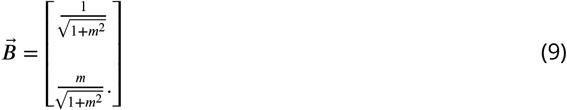
and 
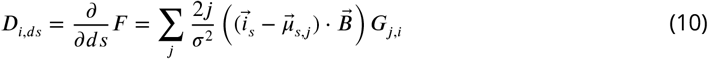

#### 3.3 Derivative with respect to *A* and *Bg*

For *A* the derivative is 6 without the *B* term and for derivative with respect to *Bg* it is unity.

## 4. Polynomial Model

For the time-lapse images, MTrack uses a polynomial model to do the fitting, which allows tracking of bending and crossing microtubules. We use a 3^*rd*^ order polynomial and modify the model presented in 2 to the following: 
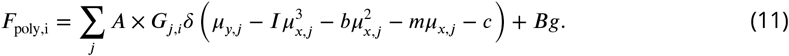

We now have two more parameters to determine *I* and *b*. *I* is the inflection of the polynomial and *b* describes the curvature of the polynomial. Now, the delta function represents putting the centroids of the Gaussians along a curve represented by the argument of the delta function. There are two more fit parameters to be determined: *b* and *I*. For doing so, we need analytical derivatives for the fit function in 1 with respect to these two new parameters. These will be derived in the following section. The spacing *ds* between the Gaussians now becomes 
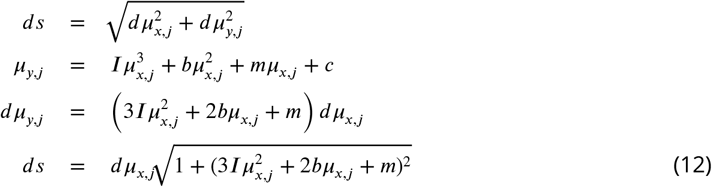

## 5. Analytical Derivatives for Polynomial Model

### 5.1 Derivative with Respect to *μ_s_, A* and *Bg*

These derivatives are the same as described for the line model.

### 5.2 Derivative with Respect to *ds*

The term *m* is now determined by the other fit parameters and can be written as 
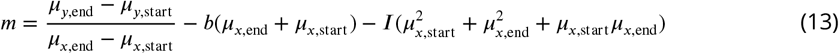

We define a new term *S* as 
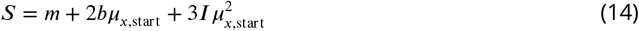

The 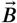 can then be redefined as 
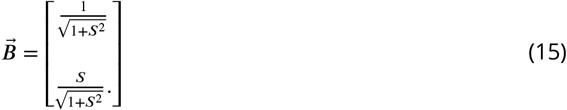

The derivative wrt *ds* can then be written as before 
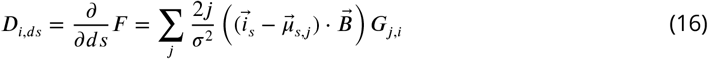

### 5.3 Derivative with Respect to Curvature (b)

We define a term *H* as 
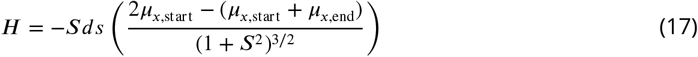

Defining a vector 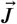 as 
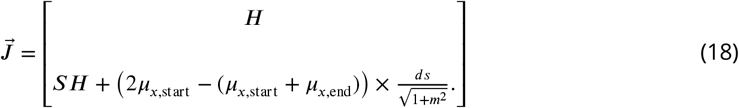

The derivative with respect to *b* can then be written as before 
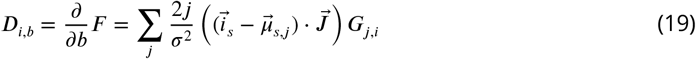

### 5.4 Derivative with Respect to Inflection (*IF*)

We define a term *K* as 
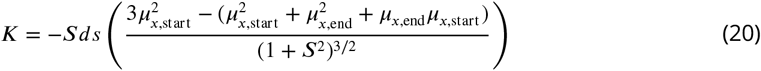

Defining a vector 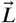 as 
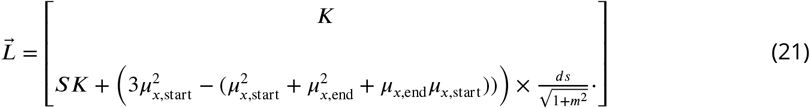

The derivative with respect to *IF* can then be written as before 
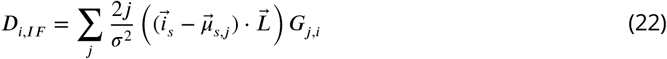

## 6. Line Parameters from MSER Ellipses

For the seed image the line parameters need to be determined. The line parameters are the slope and the intercept of the line, for a two dimensional ellipse, the covariance matrix can be represented as 
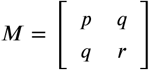

Here 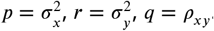. Defining a parameter dr as *dr* = 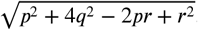. The eigenvalues of the 2 × 2 matrix are given by 
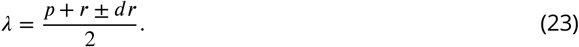

The eigenvector is given by 
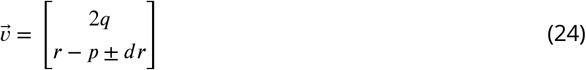

Any point (*x, y*) along the vector can be represented as 
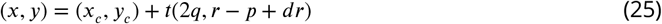

Eliminating *t* from the above vector equation gives us the line parameters (slope = m, intercept = c) of the major axis of the ellipse as 
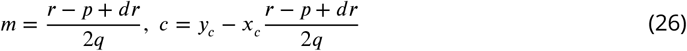

## 7. The Levenberg-Marquradt Solver

In order to obtain the optimized set of parameters, we use the Levenberg-Marquradt solver to minimize the sum of squared differences in Eq.1. The *χ* squared function can be written as 
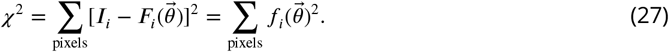

To do so, we perform Taylor expansion on the *χ*^2^ function as 
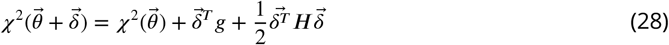
*δ* is given by 
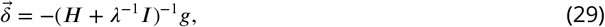
and the matrix *H* is given by 
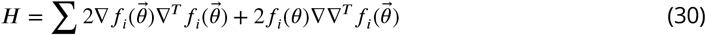

As the matrix *H* is sparse, the second term containing the 2^*nd*^ order derivative is ignored and only the first term is kept. We minimize the *χ* squared function with respect to delta and if the solution is going towards minima the *λ* parameter is decreased by a factor of 10, else increased by the same factor.

## 8. Gaussian Mask Fits

After obtaining the optimized set of parameters from the Levenberg-Marquradt solver, we refine the obtained result by doing a weighted sum of Gaussian mask fits [40] to further improve the accuracy of the detection. For this step we construct the mask for the start and end positions as 
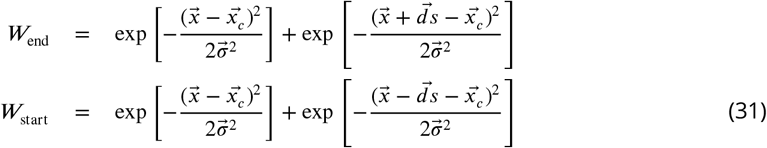

This is an iterative process of finding the location of the mask constructed by sum of two Gaussians and at the end of the iteration we obtain a sub-pixel accurate localization of the end points of the microtubule. The process is repeated for each frame and for all the detected microtubules.

## 9. One-to-one Mapping of Microtubules to MSER Regions

For seeds as well as (non-colliding) dynamic microtubules, there is a one-to-one mapping of the detected microtubules with the MSER-segmented regions. In order to make an initial guess for the start and the end position, we use the slope of the line determined for the seeds and determine the intersection point of the line with the MSER-segmented region as the initial guess for the start and the end position of the dynamic microtubule.

For the dynamic channel, we draw a line from the start and end position of the previous frame and determine the intersection point of the drawn line with the MSER segmented region, along that line we look for the intensity points which are at least greater than 50% of the maximum intensity value in that region. The guess for the polynomial function, which the microtubule growth, follows is taken from the polynomial function determined in the previous frame. For the first frame in the dynamic channel the line parameters are used from the seed image frame and the polynomial parameters are assumed to be 0 and then determined via the optimizer.

## 10. Handling Colliding Microtubules

Each microtubule evolves according to a polynomial function, whose parameters are determined in each frame using the function parameters of its evolution in the previous frame. Including prior knowledge helps making good initial guesses for the localization to proceed in the current frame. As the dynamic function of evolution of each microtubule is smooth and unique, the optimizer is unlikely to make mistakes and if it does the program can recognize that by noting sharp changes in the polynomial function parameters of growth of a given microtubule. Two special cases are discussed with respect to colliding microtubules.

### 10.1 Single MSER Region for Multiple Colliding Microtubules

In a scenario where a single segmented region contains multiple microtubules, the optimizer relies on the polynomial parameters for the microtubule determined in the previous frame and is able to correctly determine the growing end points. MTrack keeps track of the angular change in the direction of the dynamic microtubules and determines a mistake if the angular change is greater than a user defined value, which by default is 20 degrees.

### 10.2 Multiple MSER Regions for a Single Microtubule

When microtubules are close or collide, it may happen that a single microtubule finds itself in multiple MSER segmented regions. In such a case, the optimizer will fit in all regions to determine the growing end point location and will choose the point closest to the last known location of the microtubule.

## 11 Simulations

Microtubules are simulated using linear models and 3^*rd*^ order polynomials, where the parameters of each function are determined by a random number generator. Images are created by convolution with the PSF, adding a background, and rendering the final pixel intensities using a Poisson process. The simulation code is available on Github https://github.com/PreibischLab/MTrack/tree/master/src/main/java/dummyMT.

## 12 Documentation and Installation

Detailed documentation and installation instructions are available on the ImageJ wiki http://imagej.net/MicrotubuleTracker.

## 13 Code and Availability

The code is open-source, mostly implemented in ImgLib2 [29] and is provided as Fiji [36] plugin. The iterative RANSAC for functions is based on the *mpicbg* package written by Stephan Saalfeld [35]. The source code is available on GitHub https://github.com/PreibischLab/MTrack. It is released under GPLv3.

## 14 Example Data

An example dataset is available for download on the ImageJ wiki http://imagej.net/MicrotubuleTracker#Example including the MTrack parameters to successfully run and analyze the demo movie. The expected runtime is 10–20 min on standard hardware.

## Acknowledgements

We thank Tobias Pietzsch (Tomancak lab, MPI-CBG, Dresden) for implementing MSER in ImgLib2, Curtis Rueden (LOCI, UW Madison) for Fiji maintenance and support, Hadrien Mary (Brouhard lab, McGill University, Montréal) and Mohammed Mahamdeh (Howard lab, Yale University) for constructive criticism and comments on the manuscript. We thank the AMBIO imaging facility (Charité, Berlin), where the experimental data were acquired. We thank all former and current members of the Preibisch and Reber labs for discussion and helpful advice. VK thanks David Hörl (LMU München / MDC Berlin) for stimulating discussions.

VK was supported by the IRI Life Sciences postdoc fellowship in the labs of SR and SP. CH and SR acknowledge funding by the IRI Life Sciences (Humboldt-Universität zu Berlin, Excellence Initiative/DFG). WH was supported by the Alliance Berlin Canberra “Crossing Boundaries: Molecular Interactions in Malaria”, which is co-funded by a grant from the Deutsche Forschungsgemeinschaft (DFG) for the International Research Training Group (IRTG) 2290 and the Australian National University. SP was supported by the MDC Berlin.

## Author Contributions

SP and SR conceived the project. VK implemented the tracking, the GUI and developed the derivations. SP implemented the RANSAC function framework. WH and CH provided the experimental data and guided software optimizations. SP and SR supervised the project. SP and SR wrote the paper with input from VK.

## Supplementary Figures and Movies

1. Supplementary Figure 1: Flow Diagram of MTrack Detecting, Tracking, and Analyzing Dynamic Microtubules
2. Supplementary Figure 2: Microtubule Seed Detection
3. Supplementary Figure 3: Comparison between Manually Analyzed and Computer-inferred Microtubule Dynamics
4. Supplementary MovieS1: Illustration of the Component Tree

